# Genetic load proxies do not predict fitness better than inbreeding does in a wild population

**DOI:** 10.1101/2025.07.07.663445

**Authors:** D. A. Robledo-Ruiz, P. Sunnucks, B. Quin, M. J. L. Magrath, D. Harley, A. Pavlova

## Abstract

A key goal in conservation is managing endangered populations in order to maximise individual and population fitness. Genetic management that reduces inbreeding and increases genetic diversity has been consistently successful, but recent attention has been drawn to managing genetic load instead. We investigated whether genetic load proxies based on evolutionary conservation predict individual fitness better than inbreeding does. We re-sequenced the genomes of 43 wild helmeted honeyeaters with lifetime fitness data, and 28 individuals of other subspecies. None of the 28 genetic load proxies we tested predicted lifetime reproductive success better than three inbreeding metrics did, and different load proxies were inconsistent regarding the occurrence of purging or accumulation of deleterious alleles. Genetic load proxies need further validation as fitness indicators for conservation management.

## Main Text

A central question in conservation is how to manage threatened populations to improve long-term viability. Small, isolated endangered populations are at risk of extinction through inbreeding depression (i.e., lower fitness in the progeny of related individuals) and reduced evolutionary potential (*1*). The classic approach of reducing inbreeding and increasing genetic diversity—by augmenting gene flow into genetically eroded populations—is effective at alleviating inbreeding depression and improving population fitness and growth (*2*). The recommendation has been to choose large, outbred source populations that are not problematically divergent from the target population (*1*). Strong support for this approach comes from experimental studies and cases of species being brought back from the brink of extinction (*2*, *3*). Recently, however, concerns arose about possible overemphasis on genome-wide genetic diversity while overlooking the effects of harmful genetic variation (i.e., genetic load) on fitness (*4*). The argument is that gene flow from large, diverse populations would introduce harmful alleles, and that historically small- to medium-sized populations that have been genetically purged (i.e., depleted from harmful recessive alleles by inbreeding-enhanced purifying selection; *5*) should be used as sources instead (*6*, *7*). This latter approach has been systematically challenged (*8*, *9*).

Individual genetic load is the potential or actual *reduction in fitness* caused by harmful mutations in an individual’s genome, and equal to the sum of masked load (i.e., load at heterozygous sites that is not expressed) and realized (or expressed) load (*5*, *10*, *11*). However, because obtaining fitness data for wild populations is challenging, conservation genomic studies are instead increasingly focussing on assessing genome variation and the predicted harmful effects of mutations. Predicted deleteriousness scores of mutations are summarized into metrics that aim to approximate individual genetic load. These *proxies* of genetic load are then used to infer individual- or population fitness, and whether a population has undergone purging. While most tools developed to predict mutation deleteriousness were originally designed for humans and model organisms (e.g., GERP, CADD, SNPeff; *12–15*), only one has been tested against fitness in wild populations (SNPeff; *16*, *17*). Despite the growing use of genetic load proxies to guide genetic management (*17–21*), the fundamental assumption that managing genetic load will lead to better conservation outcomes—because genetic load proxies are superior fitness predictors— has yet to be thoroughly tested. Therefore, studies seeking to validate genetic load proxies should assess whether they are more effective predictors of fitness than is level of inbreeding (*22*).

Genomic Evolutionary Rate Profiling (GERP) is a tool that gives derived alleles a deleteriousness score based on the level of evolutionary conservation of the sites in which they occur: higher scores are given to derived alleles at otherwise highly conserved genomic sites (*12*, *13*). These scores have been summarized into individual genetic load proxies in diverse ways (*10*), although explanations of choices are limited (for an exception see *20*). An implicit assumption is that these proxies are interchangeable, which in practice has translated into studies reaching similar inferences (e.g., the occurrence of purging) for different organisms through different proxies.

Here, we combine genomic data from multiple subspecies of the yellow-tufted honeyeater (*Lichenostomus melanops*; *23*), and individual fitness data (*24*) for the endangered subspecies *L. m. cassidix*, to test the key assumption that, for wild populations, genetic load proxies predict individual fitness better than inbreeding metrics do. By applying different GERP-based proxies from eight published studies (Table 1), we also explore whether different load proxies have the same power for predicting fitness, and whether they lead to consistent conclusions about purging versus accumulation of genetic load.

**Table 1.** Summary of the studies and genetic load proxy calculations applied in this study. The last row contains three new proxies assessed in this study. Proxies marked with * were those not calculated by the published studies but were derived here for comparison purposes.

| Study | Species | Compared to historic/relative | Type | Genetic load proxies | GERP score threshold |
| --- | --- | --- | --- | --- | --- |
| Grossen et al. (2020) *Nat. Comms.* | Alpine ibex | Relative (Iberian, Nubian and Siberian ibex, Markhor, Bezoar, domestic goat) | Count | Total load = count of derived alleles<br>Realised load = count of sites with derived alleles in homozygous state | -2-2, 2-4, 4-6, >6 |
| Mooney et al. (2023) *Mol. Biol. Evol.* | Ethiopian wolf | Relative (Arctic and Isle Royale wolf, pug, labrador retriever, Tibetan mastiff, border collie) | Count | Total load = count of derived alleles<br>Total load = count of sites with derived alleles<br>Realised load = count of sites with derived alleles in homozygous state | > 4 |
| Gonçalves-Dias et al. (2023) *Mol. Biol. Evol.* | Domesticated amaranthus | Relative (wild amaranthus) | Sum | Total load = summation of count of derived alleles weighed by their score<br>*Realised load = summation of count of sites with derived alleles in homozygous state weighed by their score | > 0 |
| Wu et al. (2023)<br>*Cell* | Potato | Relative (potato landraces) | Sum | Total load = summation of count of derived alleles weighed by their score<br>Realised load = summation of count of sites with derived alleles in homozygous state weighed by their score<br>Masked load = summation of count of sites with derived alleles in heterozygous state weighed by their score | > 2.75 |
| Dussex et al. (2021)<br>*Cell Genomics* | Kākāpō | Historical | Average | Total load = summation of count of derived alleles in homozygous state weighed by their score, divided by count of all *derived* alleles<br>*Realised load = summation of count of derived alleles weighed by their score, divided by count of all *derived* alleles | > 2 |
| von Seth et al. (2021) *Nat. Comms.* | Sumatran rhino | Historical | Average | Total load = summation of count of derived alleles weighed by their score, divided by count of all *derived* alleles | > 4 |
|  |  |  |  | *Realised load = summation of count of derived alleles in homozygous state weighed by their score, divided by count of all *derived* alleles |  |
| Kutschera et al.<br>(2022) *BMC Bioinformatics* | Sumatran rhino | Historical | Average | Total load = summation of count of derived alleles weighed by their score, divided by count of *derived* alleles at 99th percentile<br>*Realised load = summation of count of derived alleles in homozygous state weighed by their score, divided by count of *derived* alleles at 99th percentile | 99th percentile |
| Smeds & Ellegren<br>(2023) *Mol. Ecol.* | Scandinavian wolf | Relative (Finland and Russian populations) | Average | Realised load = summation of count of derived alleles in homozygous state weighed by their score, divided by count of all *called* alleles<br>Masked load = summation of count of derived alleles in heterozygous state weighed by their score, divided by count of all *called* alleles<br>*Total load = summation of count of derived alleles weighed by their score, divided by count of all *called* alleles | > 4 |
| This study | Helmeted<br>honeyeater | Relative (other yellow-tufted<br>honeyeater subspecies) | Average | Total load = summation of count of derived alleles weighed by their score, divided by count of all *called* alleles<br>Realised load = summation of count of derived alleles in homozygous state weighed by their score, divided by count of all *called* alleles<br>Masked load = summation of count of derived alleles in heterozygous state weighed by their score, divided by count of all *called* alleles | > 0 |

To obtain a species-wide picture of genomic variation, we re-sequenced genomes of all four recognized subspecies of *L. melanops* (mean read depth = 23.5, SD = 4.1, range = 15.8–31.8; table S1)—*L. m. cassidix* (*n* = 43), *L. m. gippslandicus* (*n* = 17), *L. m. melanops* (*n* = 5), and *L. m. meltoni* (*n* = 5)—and of *cassidix*×*gippslandicus* hybrids (*n* = 2; here after *cassidix*, *gippslandicus*, *melanops*, *meltoni* and *F1*, respectively; Fig. 1A). Subspecies *cassidix*—the helmeted honeyeater—is Critically Endangered having been reduced to a single remnant population of ∼50 individuals (*25*, *26*). Genomic inbreeding estimates and lifetime fitness data for *cassidix* individuals showed strong inbreeding depression (*23*). As previously reported, we found three genetically distinct groups: *cassidix*, *gippslandicus*, and *melanops+meltoni* (Fig. 1B and C; *27*). Subspecies *gippslandicus* is the closest relative to *cassidix* and the most genetically diverse (Fig. 1C). We found that *cassidix* experienced a population decline that started ∼61 generations ago (Fig. 1D), which coincides with the clearing of swamps by non-Indigenous human settlers in Victoria from the beginning of the 19th century (*28*). Our analyses showed that the genetic diversity of *cassidix* has been declining over time (π_cassidix_pre2000_ = 0.0069, π_cassidix_post2005_ = 0.0066; also *29*).

**Fig. 1.**
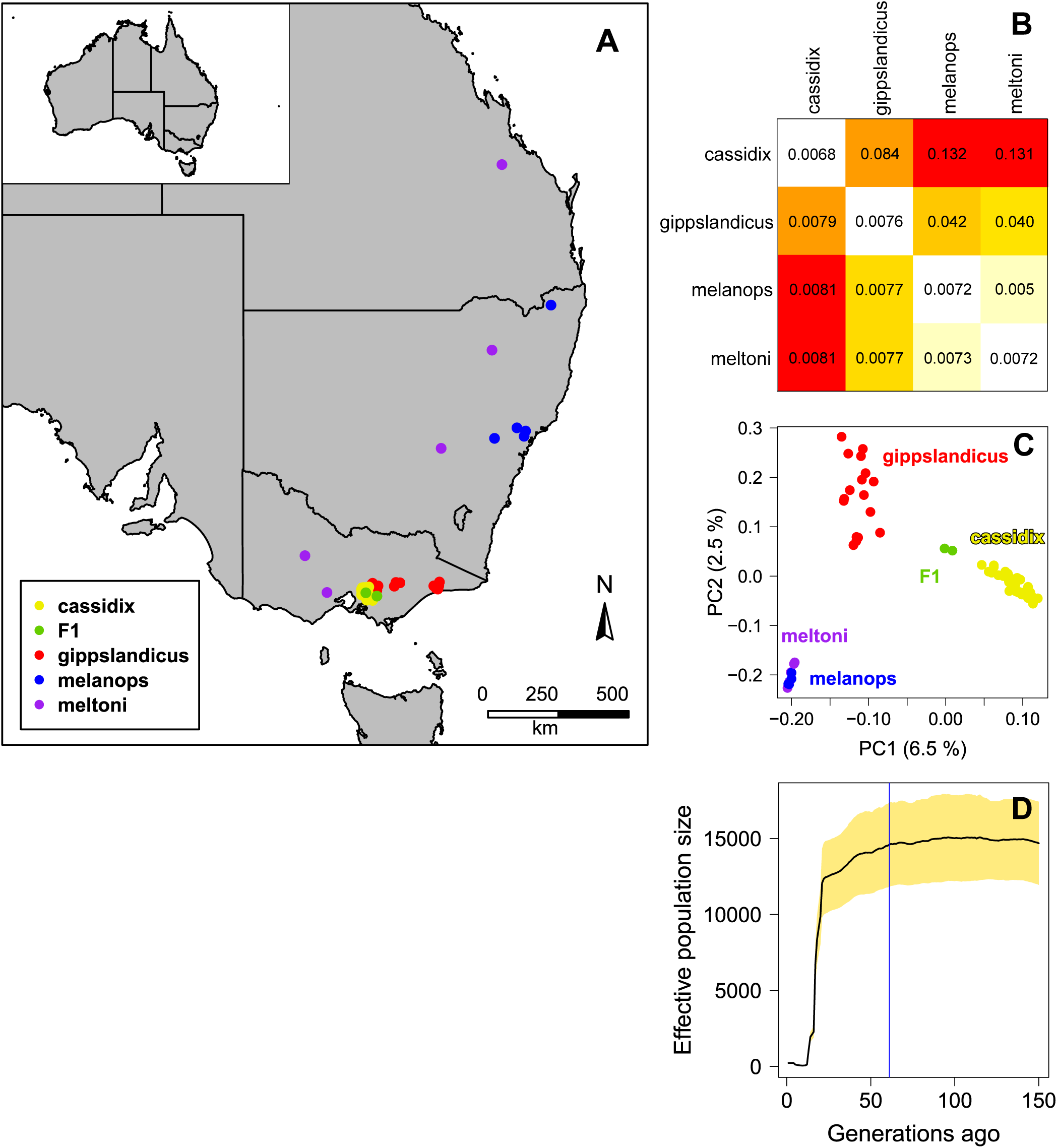
Genetic structure, diversity and demographic history of the yellow-tufted honeyeater subspecies. (**A**) Geographic distribution of the sampled individuals of the four subspecies of *Lichenostomus melanops*. Sample sizes: *n_cassidix_* = 43*, n_gippslandicus_* = 17, *n_melanops_* = 5, *n_meltoni_* = 5, *n_F1_* = 2. (**B**) Heatmap and values of differentiation (*F*_ST_; above the diagonal) and sequence divergence (D_xy_; below the diagonal) between the subspecies. Population nucleotide diversity (*π*) is included in the diagonal. (**C**) Principal component analysis of all yellow-tufted honeyeater individuals included in this study. Percentage of variance explained by each component in parentheses. (**D**) Demographic history of *cassidix* (*n* = 13). Yellow area represents 95% CI from 100 replicates of 700,000 SNPs each. The blue vertical line marks the start of the monotonic population decline 61 generations ago.

In order to assess whether genetic load proxies predict fitness better than inbreeding does, we explored the association of two individual lifetime fitness traits with 28 GERP-based genetic load proxies (Table 1) and eight inbreeding metrics. The fitness trait estimates analysed for *cassidix* individuals (subset of *23*) were lifetime reproductive success (LRS; number of fledglings produced over an individual’s lifetime, *n* = 42), and lifespan (number of days alive, *n* = 43). All genetic load proxies were calculated from the same dataset of derived alleles and their respective GERP scores. The ways in which these individual proxies differed include whether they (i) summarized realized, masked or total load, (ii) counted *alleles* or *sites*, (iii) were counts, sums of GERP scores, or average GERP scores, (iv) used the total number of *derived* alleles or *called* alleles per individual to calculate average GERP scores; and (v) what GERP scores were used as thresholds to consider derived alleles deleterious (Table 1; *20*, *21*, *30–35*). We measured individual inbreeding as the proportion of the genome in Runs of Homozygosity (ROHs, homozygous chromosome stretches ≥100 SNPs caused by matings between relatives) and as genome-wide homozygosity. We used a likelihood-based method to identify identity-by-descent ROHs, while accounting for sequencing error and mutation rates (*36*). Seven different ROH inbreeding coefficients were calculated per individual: four based on ROHs length (F_ROH(3Mb+)_, F_ROH(short)_, F_ROH(medium)_, F_ROH(long)_), two based on ROHs that arose before versus during *cassidix*’s population decline (i.e., starting ∼61 generations ago; F_ROH(old)_ and F_ROH(recent)_, respectively), and one including all detected ROHs (F_ROH_). We calculated these inbreeding coefficients because they are predicted to have different associations with inbreeding depression. For example, shorter ROHs (e.g. F_ROH(short)_, F_ROH(medium)_) signal inbreeding of ancestors in deep history (i.e., after enough time has passed for long ROHs to be broken up into smaller ROHs) and are likely less important for inbreeding depression because some of their deleterious alleles may have been purged over time, or otherwise compensated for by natural selection (*36*, *37*). On the other hand, ROHs that arose during *cassidix*’s population decline (F_ROH(recent)_) capture inbreeding due to anthropogenic causes. Finally, we calculated genome-wide homozygosity (F_HOM_), a common straightforward proxy for individual inbreeding previously used to quantify inbreeding depression in *cassidix* (*23*). The relationship of each genetic load proxy and inbreeding metric to each fitness trait was assessed with generalised linear models. We controlled for sex because there are differences in the severity of inbreeding depression between male and female helmeted honeyeaters (*23*). *P*-values were adjusted for multiple testing. To limit the number of tests being adjusted for, the number of genetic load proxies used was restricted to a subset in recent literature (Table 1).

Although proxies of realised load predicted individual LRS to some degree (*n* = 42, *R^2^* range = 0.19–0.298, *adjusted p* < 0.001), they were not the best predictors (Fig. 2 and S3). Three inbreeding coefficients predicted LRS better than did any of the genetic load proxies tested: F_ROH(3Mb+)_, F_ROH(long)_, and F_ROH_ (*R^2^_F(ROH 3Mb+)_* = 0.361, *R^2^_F(ROH long)_* = 0.316, *R^2^_F(ROH)_* = 0.313, respectively). The LRS model with the best inbreeding coefficient predictor, F_ROH(3Mb+)_, yielded an *R^2^* 21.1% higher than the best genetic load proxy, Grossen’s count of sites homozygous for derived alleles of “extreme” deleterious effects (i.e., GERP > 6, *R^2^_Grossen6+_* = 0.298; hereafter ‘*realised Grossen 6+*’). In addition, LRS models with F_ROH(3Mb+)_, F_ROH(long)_ and F_ROH(recent)_ as predictors had narrower 95% CIs than all of the genetic load proxy models. The most rudimentary estimate of inbreeding—F_HOM_—predicted LRS as effectively, or almost so, as genetic load proxies calculated as averages, but worse than count and sum proxies.

**Fig. 2.**
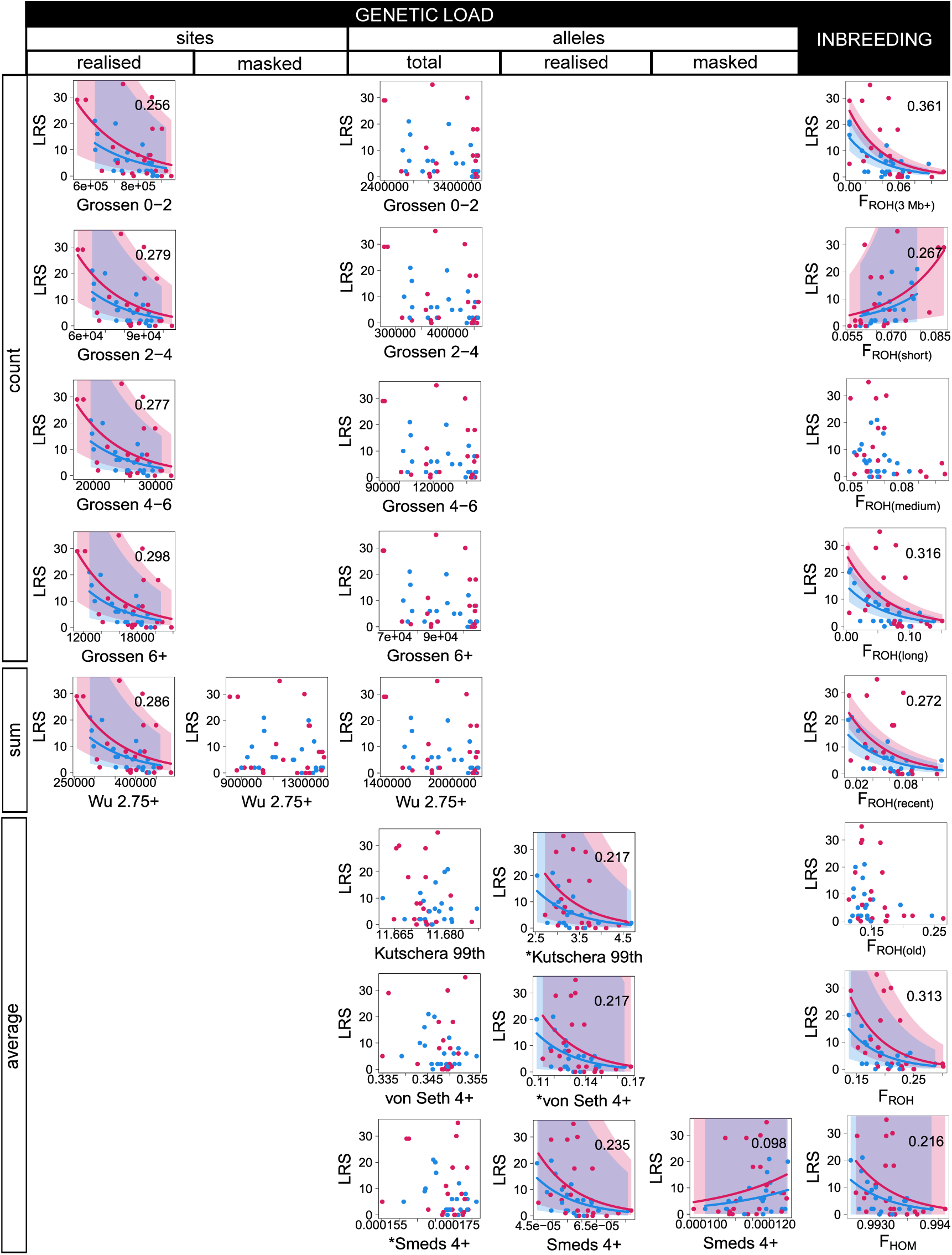
Lifetime Reproductive Success (LRS, number of fledglings produced over an individual’s lifespan; y-axes) and its relationship to proxies of individual genetic load (GERP scores; x-axes) and to inbreeding coefficients (*F*; x-axes). Genetic load proxies are grouped by columns according to whether they use counts of derived *alleles* or the *sites* in which derived alleles occurred, and whether they represent total, realised or masked load. Grouping by rows follows whether the proxies were counts of derived alleles/sites, the sum of the GERP scores for the derived alleles/sites, or average GERP scores. Names of genetic load proxies include the GERP threshold above which a mutation was considered deleterious; asterisks indicate proxies that were not calculated in the original publications but were calculated in this study for comparison purposes. Each dot represents an individual (females in red, males in blue). Trend lines represent statistically significant generalised linear models (*multiple-testing adjusted p* < 0.001), and shaded areas 95% CI. *R^2^* values are placed on the top right corner of statistically significant models.

Surprisingly, Grossen’s counts of sites with “moderately” deleterious derived alleles (*R^2^_Grossen2-4_* = 0.279) predicted LRS to the same degree as those with “highly” deleterious derived alleles (*R^2^_Grossen4-6_* = 0.277; Fig. 2 and S3). Similarly, despite most average proxies focusing on derived alleles with “moderate” to “extreme” effects (i.e., GERP > 2; *R^2^_Kutschera99th_* = 0.217, *R^2^_Dussex2+_* = 0.215, *R^2^_vonSeth4+_* = 0.217, *R^2^_Smeds4+_* = 0.235), they performed worse than the count proxy that focused on “neutral or near neutral” derived alleles (*R^2^_Grossen0-2_* = 0.256). These two findings are unexpected if higher estimated levels of deleteriousness (i.e., larger GERP scores) accurately reflect greater negative impacts on fitness. To the extent that GERP scores reflected LRS, these results caution against ignoring putatively less-harmful derived alleles, as some studies did (e.g., *20*, *34*).

Except for F_ROH(short)_—the best predictor of individual lifespan (*n* = 43, *R^2^_F(ROH short)_* = 0.099, *adjusted p* < 0.001)—overall, realised load proxies predicted lifespan better than inbreeding did (Fig. 3 and S4). However, the predictive power of realised load proxies for lifespan was much lower (*R^2^* range = 0.022–0.072) than that of inbreeding metrics for LRS (*R^2^* range = 0.216– 0.361). Given that LRS encompasses survival and reproduction at all life stages, the total effects of inbreeding depression are necessarily captured better by LRS than by lifespan (*23*, *38*, *39*).

**Fig. 3.**
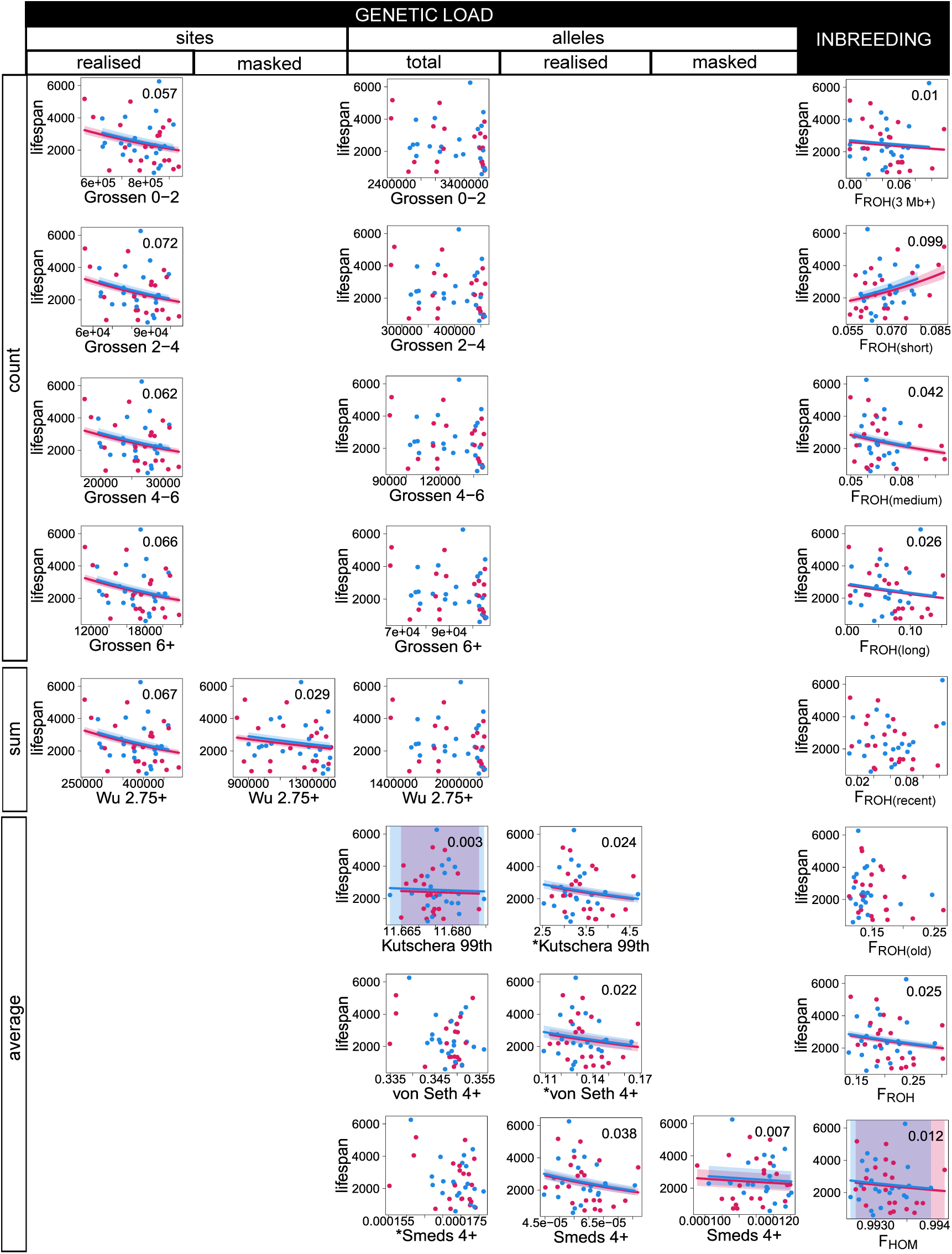
Lifespan (number of days alive; y-axes) and its relationship with proxies of individual genetic load (GERP scores; x-axes) and to inbreeding coefficients (*F*; x-axes). Genetic load proxies are grouped by columns according to whether they use counts of derived *alleles* or the *sites* in which derived alleles occurred, and whether they represent total, realised or masked load. Grouping by rows follows whether the proxies were counts of derived alleles/sites, the sum of the GERP scores for the derived alleles/sites, or average GERP scores. Names of genetic load proxies include the GERP threshold above which a mutation was considered deleterious; asterisks indicate proxies that were not calculated in the original publications but were calculated in this study for comparison purposes. Each dot represents an individual (females in red, males in blue). Trend lines represent statistically significant generalised linear models.

Additionally, LRS is more important than lifespan for the evolutionary trajectory and persistence of a population because LRS determines the genetic composition of new generations.

We next investigated whether inbreeding *and* genetic load predicted LRS better than inbreeding alone, or in other words, whether the best genetic load proxy (*realised Grossen 6+*) predicted a portion of LRS that the best inbreeding coefficient (F_ROH(3Mb+)_) did not. After accounting for the statistical effects of sex and F_ROH(3Mb+)_ on LRS, the statistical effect of *realised Grossen 6+* was not significant nor did it improve the prediction of LRS (*p* = 0.06). Given the strong positive correlation between F_ROH(3Mb+)_ and *realised Grossen 6+* (*r*(_40_) = 0.753, *t* = 7.23, *p* < 0.001), we also explored whether combining them into the variable that maximally predicted LRS would improve the prediction of LRS overall; this combined variable increased prediction of LRS (*R^2^*) by only 0.4%.

In addition to predicting fitness, genetic load estimates have been used to determine whether populations that have declined in size—and experienced higher inbreeding—have been purged of some of their deleterious alleles. This is tested by comparing the genetic load of the focal population to that of (i) a historical or ancient sample, or (ii) a related population that is larger or has a demographic history less marked by decline. Lower total (i.e., realised + masked) genetic load in the focal population has been interpreted as evidence of purging, while the opposite suggests the accumulation of deleterious alleles (*21*, *35*, *34*). In order to explore whether all total load proxies yielded the same inference (i.e., purging or accumulation of putatively harmful mutations), we compared the genetic load proxies of the most recent *cassidix* individuals (i.e., born after 2005, *n* = 17; hereafter *cassidix post2005*) to those of (i) *cassidix* born before 2000 (*n* = 26; hereafter *cassidix pre2000*); and (ii) the relatively non-bottlenecked *gippslandicus* and *melanops*+*meltoni*.

Total load proxies gave conflicting signals of whether *cassidix post2005* experienced purging of putatively harmful genetic variation relative to that of *gippslandicus*. Of the seven total load proxies for which significant differences between *cassidix post2005* and *gippslandicus* were found, count and sum proxies indicated purging (*n* = 4), while average proxies indicated accumulation of deleterious alleles (*n* = 3; Fig. 4, S5 and S6). The remaining total load proxies showed no significant differences (*n* = 5). It was notable that Mooney’s two proxies of total load, which differ only in whether they count derived *alleles* or the *sites* at which derived alleles occurred, pointed towards different conclusions: no significant differences and purging, respectively. Comparison of the historical sample (*cassidix pre2000*) and *cassidix post2005* showed no significant differences for any total load proxy, despite the presence of strong inbreeding depression (*23*).

**Fig. 4.**
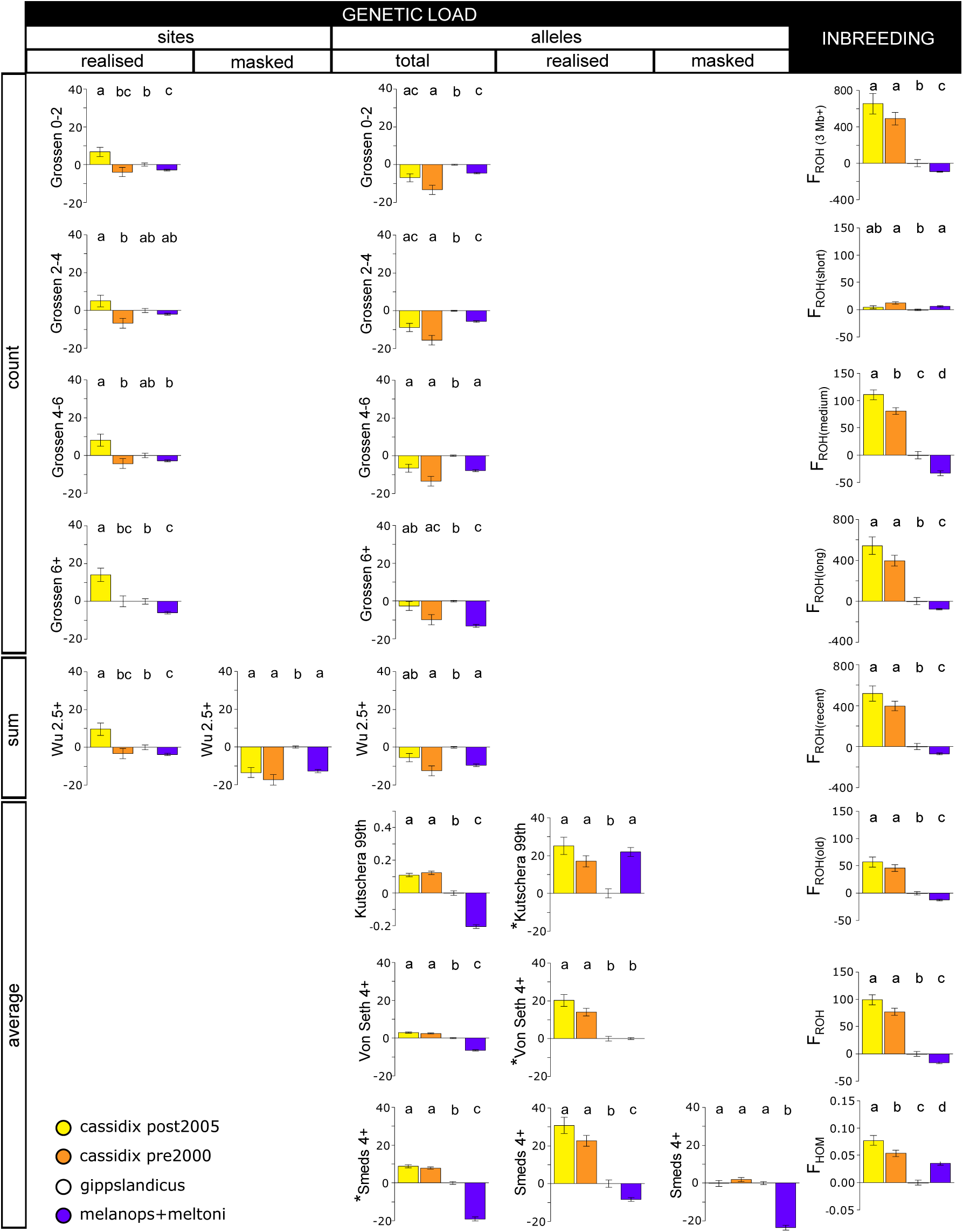
Relative differences in proxies of individual genetic load (GERP scores) and inbreeding coefficients (*F*) between yellow-tufted honeyeater genetic groups and time periods and time periods (*cassidix post2005, cassidix pre2000*, *gippslandicus*, and *melanops+meltoni*). Percentual differences for each group are calculated with respect to *gippslandicus*. Genetic load proxies are grouped by columns according to whether they use counts of derived *alleles* or the *sites* in which derived alleles occurred, and whether they represent total, realised or masked load. Grouping by rows follows whether the proxies were counts of derived alleles/sites, the sum of the GERP scores for the derived alleles/sites, or average GERP scores. Names of genetic load proxies include the GERP threshold above which a mutation was considered deleterious; asterisks indicate proxies that were not calculated in the original publications but were calculated in this study for comparison purposes. Bars represent mean values, and whiskers standard errors. Statically significant pairwise differences between genetic groups are represented by letters (*post hoc* Welch’s t-tests, *multiple-testing adjusted p* < 0.05).

Besides showing that GERP load proxies were not better predictors of fitness than were inbreeding metrics, our results also show that power to predict fitness varies with choice of genetic load proxy (i.e., mathematical formula and GERP thresholds; Table 1). Although inbreeding coefficients calculated from ROH also require selection of thresholds, the higher complexity of GERP introduces sources of variation not present in F_ROH_, for example, the evolutionary depth and number of species used for the input phylogeny, the units of the branches in the phylogenetic tree, the number of outgroups used to infer ancestral/derived alleles (table S3), and the criteria used to infer the ancestral/derived alleles (table S3). Importantly, the absolute value of GERP scores depends on a phylogenetic tree adequately scaled in substitutions-per-site units; if the tree is not correctly scaled, the thresholds used to identify categories of putative deleteriousness (e.g., GERP score > 6 as “extremely” deleterious; *13*) are incomparable across studies. The added complexity likely applies to other genetic load tools that rely on genome annotations rather than evolutionary conservation (e.g., CADD, SNPeff). For example, annotations differ in quality and depend on the organism for which they were developed (some are lifted over from annotations of model organisms). All these factors make findings across genetic load studies difficult to compare, even in human research (*10*).

Accumulation or purging of harmful variants has been inferred using genetic load tools in a variety of taxa (*11*). Given our results, it is unclear whether these inferences are biologically meaningful or just reflect different ways of calculating genetic load proxies. It has been noted that genetic load estimation will depend on the genomic dataset being used (e.g., the phylogeny used for inferring evolutionary conservation; *11*). Here, we demonstrate that opposite conclusions about purging can be drawn from the same genetic dataset examined via different commonly applied metrics. Our results suggest that, until genetic load tools are validated and best practices are established, caution is warranted when drawing conclusions about purging from genetic load proxies in genome data.

Very few studies have tested links between individual fitness and genetic load proxies in wild populations, and it is unknown whether annotation-based genetic load tools (e.g., SNPeff, CADD) perform better than GERP and F_ROH_. A study on the arctic fox (*Vulpes lagopus*) found a negative association between the proportion of homozygous loss-of-function (LOF) alleles and individual LRS and lifespan, which led the authors to infer that this portion of genetic load “influences” fitness (*16*). However, individual inbreeding F_ROH_ as a predictor of fitness was *not* included in their models (only expected genome-wide heterozygosity was tested as an independent predictor). Another study tested the correlations of individual inbreeding (F_ROH_ for ROH > 1Mb) and LOF genetic load (identified with SNPeff) with LRS for the northern elephant seal (*Mirounga angustirostris*; *17*). Their results were consistent with ours: F_ROH_ predicted female lifetime reproductive success *better* than LOF allele metrics did. It is yet unclear under which circumstances and for which species genetic load proxies might be better fitness predictors than F_ROH_. However, genetic load proxies are often used *as if superior to inbreeding measures*, despite little to no validation.

It is important to note that deleterious alleles within ROHs affect fitness, but power to detect them varies. For example, a study of the hihi (*Notiomystis cincta*) found that F_ROH_ was not a significant predictor of LRS, whereas 13 SNPs located within ROHs were (*40*). These loci were found with a Genome-Wide Association Study, which is a top-down method: individual fitness data are used to identify genetic variants associated with it. Genetic load proxies are used bottom-up in the *absence* of fitness data: examining genetic variants to predict individual fitness.

Several limitations of genetic load proxies have been previously pointed out, including GERP’s assumption that the strength of purifying selection is constant over time, its low precision for characterising nearly neutral mutations, and the assumption underlying annotation-based methods that biochemical effects of nonsynonymous mutations translate into fitness consequences (*41*). So far, there is no consistent evidence to support genetic load proxies (as they have been implemented) as the superior fitness predictors, nor that they should replace inbreeding coefficients to guide conservation programs. Wildlife managers should be fully informed about the assumptions and uncertainties surrounding genetic load proxies. Conservation programmes where proxies have been assumed to be the best predictor of fitness, or purging has been inferred, should be revisited.

As more genomic data become available, more studies from wild populations with lifetime fitness data are needed to validate and benchmark different genetic load proxies. Improvements in the power of genetic load proxies to predict fitness would be a positive step forward, opening exciting research avenues. However, even if perfected tools allowed the management of populations to minimize number of harmful alleles and maximize fitness, retaining genetic variation would still be needed to maximise long-term population viability (*8*, *28*, *42–44*). This includes retention of harmful alleles, as some of them may underpin future adaptations (*45*). Preservation of evolutionary potential without compromising current fitness can be achieved by masking the effects of harmful recessive alleles (*46*) through managing for low levels of inbreeding.

## Acknowledgments

We acknowledge the First Nations throughout Australia, recognise their continuing connection to land, waters and culture and pay our respects to their Elders past, present and emerging. This research was conducted on Wurundjeri Woi-wurrung Countries. We thank Y. P. Lee, R. Turakulov, M. Kardos, and J. Castrejón-Figueroa for technical assistance, and C. van Oosterhout for helpful comments.

## Author contributions

Conceptualization: PS, DARR, AP

Methodology: DARR, PS, AP, BQ, MJLM

Formal analysis: DARR

Investigation: BQ, MJLM, DH

Data curation: BQ, MJLM, DARR

Writing – original draft: DARR

Writing – review & editing: DARR, PS, AP, BQ, MJLM, DH

Visualization: DARR

Supervision: PS, AP, MJLM, DH

Project administration: AP, PS, MJLM, DH

Funding acquisition: AP, PS, MJLM, DH, DARR

## Supplementary Materials

### Materials and Methods

Figs. S1 to S6

Tables S1 to S3

