## Supplementary Materials for "Genetic load proxies do not predict fitness better than inbreeding does in a wild population"

**The PDF file includes:**

Materials and Methods
Figs. S1 to S6
Tables S1 to S3

### Materials and Methods

Sample collection and sequencing

We used existing DNA samples extracted from the blood of 45 *cassidix*, 18 *gippslandicus*, 2 *cassidix*×*gippslandicus* hybrid (hereafter '*F1*'), 5 *melanops* and 5 *meltoni* individuals that had previously been examined for genetic variation using DArT SNP data, obtained using DArTseq, a reduced representation sequencing technology (*23*, *24*). Samples were collected between 1989 and 2012 in Victoria and New South Wales, Australia. Lifetime fitness data were available for the *cassidix* individuals: lifetime reproductive success (LRS) was measured as the total number of fledglings produced by an individual in its lifetime, and lifespan was measured in number of days (*24*).

Genomic DNA library construction and sequencing were performed by the Deakin University Genomics Centre. Libraries were quantified with Qubit Broad-range and High-sensitivity assays (Invitrogen, USA) using a Qubit 3.0 Fluorometer (Invitrogen, USA). For each sample, 100–250 ng of genomic DNA (in a volume of 52.5 µL) was sonicated to target size of 350 bp using a Q800R sonicator (Qsonica, CT, USA) with the following parameters: 20% amplitude, pulse on 15 seconds, pulse off 15 seconds, temperature 4°C for 10 minutes. The sonicated DNA was purified with 1X volume of AMPure XP magnetic beads (Beckman Coulter, Indianapolis, IN), and then per sample sequencing libraries were prepared using the NEBNext Ultra DNA Library Prep Kit for Illumina (New England Biolabs, Ipswich, MA) according to the manufacturer's protocol. In the adaptor ligation step, the NEBNext adaptor for Illumina (provided at 15 µM) was diluted 10-fold in dilution buffer (10 mM Tris-HCI, 10 mM NaCl) (Astral Scientific, Australia). The PCR enrichment of adaptor-ligated DNA cycling conditions, denaturation, and annealing/extension cycle steps were repeated with a total of 8 cycles. Quantification and size estimation of the libraries was performed on the Qubit 3.0 Fluorometer and 4200 TapeStation System (Agilent, Santa Clara, CA).

Two µL of each library were pooled into a new microfuge tube and enzymatically treated with Illumina Free Adapter Blocking Reagent (Illumina, San Diego, CA). The pooled library was pre-sequenced on the MiniSeq Sequencer (2 × 150 bp paired-end reads) (Illumina, San Diego, CA) to obtain the read distribution of each sample. Each library was then re-pooled to equal molar concentrations, enzymatic treated, denatured and normalised to 2 nM. Finally, the library was sequenced on the NovaSeq 6000 Sequencer (2 × 150 bp paired-end reads) (Illumina, San Diego, CA). Overall, sequence data were generated for 75 individuals with an average sequencing depth of 22.9× per sample (table S1).

Genotyping

*Genotype calling.* Genotypes were called using a containerized genotype-calling pipeline based on GATK (trust1/gatk v4.1.4.1, available at: https://hub.docker.com/r/trust1/gatk). Briefly, raw Illumina reads for 75 individuals were mapped to the helmeted honeyeater chromosome-length genome (GenBank accession VLJF00000000.2; *23*) with BWA MEM v0.7.17 (*47*), and alignment files were sorted and deduplicated. Variants per sample were pre-called with HaplotypeCaller, followed by a first and second pass of base-quality score recalibration with BaseRecalibrator and ApplyBQRS, according to GATK best practices. A second round of HaplotypeCaller was performed on the recalibrated bam files, and GenomicsDBImport was used to merge the multiple GVCFs. For the final step, due to a bug in GATK v4.1.4.1 GenotypeGVCFs, we used GATK v4.1.5.0 for joint genotyping of all 75 samples with argument --include-non-variant-sites set to 'true' in order to obtain both variant (polymorphic) and invariant (monomorphic) sites. We retained invariant sites in order to calculate unbiased population nucleotide diversity ($\pi$), sequence divergence ($D_{xy}$) and individual heterozygosity estimates (see below; *48*).

*Filtering.* VariantFiltration in GATK was used to carry out hard filtering of variant sites with QD < 2.0, MQ < 40.0, FS > 60.0, ReadPosRankSum < −8.0, MQRankSum < −12.5, and SOR > 3.0. The software vcftools (*49*) was used to remove loci within previously identified repetitive regions (*23*), sex chromosomes, mitogenome, indels, and sites with higher than twice and lower than 1/3 the average read depth across individuals (i.e., 7.91 and 47.4 read depth, respectively). Genotypes (i.e., genotype of an individual at a specific locus) with meanDP < 5 were set as missing. In order to reduce genotyping errors included in the dataset, we removed sites missing in more than 50% of samples, or with MAC < 3 (i.e., guaranteeing that each variant is present in at least two individuals). Aiming to exclude sites erroneously fused during genome assembly (i.e., 'multilocus' SNPs, *50*), we removed *excessively heterozygous* sites according to HW-equilibrium expectations. Three of the 75 original samples (2 *cassidix* and 1 *gippslandicus*) were excluded from downstream analyses due to suspected sample contamination. The filtered dataset containing invariant and variant sites consisted of 847,367,083 sites (hereafter 'Dataset 1'). After the removal of invariant and multiallelic sites, 26,527,283 biallelic SNPs were retained to create a second dataset (hereafter 'Dataset 2').

<u>Subspecies differentiation analyses</u>

*Unbiased $\pi$, $D_{xy}$, $F_{ST}$ and individual heterozygosity.* Dataset 1 was used as input for the software *piawka*, which calculates unbiased per-site summary statistics from sequence data containing invariant and variant sites, including multi-allelic ones (*51*). The software *piawka* was inspired by *pixy* (*48*) which calculates average heterozygosity per *called site*. However, unlike *pixy*, *piawka* includes multiallelic sites whose removal can bias diversity estimates (*52*). We used *piawka* to calculate population nucleotide diversity ($\pi$) for each subspecies, and between-population sequence divergence ($D_{xy}$) and fixation index ($F_{ST}$) for each subspecies pair. We also calculated $\pi$ for the most recent *cassidix* individuals (i.e., born after 2005, $n$ = 17; hereafter *cassidix_post2005*) and *cassidix* born before 2000 ($n$ = 26; hereafter *cassidix_pre2000*).

*PCA.* Dataset 2 was filtered in order to remove sites with any missing data. Linkage Disequilibrium pruning (LD; 50 kb window size, 10 kb step size, $r^2 = 0.5$; see *53*) and a Principal Component Analysis (PCA) were performed with plink v1.9 (*54*).

<u>Demographic history</u>

We used the LD-based method implemented in GONE (*55*) to estimate the change in the effective population size ($N_e$) of *cassidix* up to 150 generations ago. We used Dataset 2 and kept only SNPs present in the 25 autosomes for which a high-density linkage map was available (*23*). Regions of chromosomes 13 and 26 that were likely to be misassembled (*23*) were excluded from this analysis. We used genetic data of 13 *cassidix* birds with hatch dates between 1989 and

1991 in order to analyse one cohort. This sample represents ~19% of the wild population whose size was ~70 individuals between 1989 and 1991. We kept SNPs present in 100% of individuals and with MAF 0.08. We created 100 subsets of 700,000 SNPs chosen at random (i.e., 100 replicates). The genetic distance of each SNP was interpolated by adjusting a LOESS curve (span = 0.2) to the linkage map. GONE was run for each of the 100 SNP subsets allowing only 50,000 SNPs per chromosome ($c$ = 0.05), and replicates were used to calculate confidence intervals. Historical $N_e$ was inferred assuming a generation time of 3.17 years (*56*).

Demographic history reconstruction was not possible for *gippslandicus,* whose samples did not represent a coherent population (due to being collected across multiple geographic locations and in different years). We ran GONE on three different subsets of *gippslandicus* individuals: (i) all 17 individuals, (ii) eight birds collected from the same small geographical region (i.e., Upper Yarra ranges, VIC), and (iii) a random sample of 13 individuals to match the sample size used for *cassidix*. The three subsets produced widely different demographic histories, and due to the paucity of information about the samples—including hatch dates—we could not discern which subsets violated the assumptions of GONE (i.e., absence of population structure and non-overlapping generations; fig. S1).

#### GERP

Genomic Evolutionary Rate Profiling (GERP) is an approach to identify the strength of purifying selection on putatively functional genomic regions by quantifying their substitution rate and comparing it to the expected neutral rate (*12*). By using a multiple sequence alignment of different species genomes and a phylogenetic tree, GERP++ assigns a substitution deficit score to each nucleotide position, whose magnitude captures the number of substitutions "rejected" by evolutionary constraint at that position (i.e., higher scores reflect stronger purifying selection; *13*). To identify genomic regions under strong evolutionary constraint in the yellow-tufted honeyeater, we followed a slightly modified version of the pipeline GenErode (*34*). We used 151 publicly available avian genomes selected based on their coverage and assembly level, and aiming to proportionally represent all avian orders, suborders and families (table S2). We used only avian species for the phylogeny in an attempt to closely follow the methods of most published studies followed as reference (table S3). TimeTree was used to obtain a dated phylogenetic tree of the 151 selected birds and the yellow-tufted honeyeater, as needed by GERP++ (fig. S2; *57*).

A multi-species alignment was created by converting each of the 151 genomes to 50 bp reads with BBMap reformat.sh (*58*), and aligning them to the *cassidix* genome with BWAmem v0.7.17 (*47*). Next, we converted the alignment BAM file for each species to FASTA using a combination of samtools *mpileup* and two python scripts modified from the GenErode pipeline (*59*). All FASTA files were concatenated to create the input multi-species alignment file for GERP++. *gerpcol* was then used to calculate conservation scores for each site in the *cassidix* genome for which at least three of the 151 species were aligned. We used parameter -s to specify an evolutionary rate of 0.002 substitutions per site per million years (i.e., Aves estimate; *60*), and parameter -j to project out the reference genome in order to avoid biases (*20*). Only GERP scores > 0 were used for downstream analyses (hereafter 'valid GERP scores') because (i) GERP++ assigns 0 to positions for which fewer than 3 species were aligned (GERP++ manual), which confounds null results with authentic 0 scores, and (ii) negative scores are not informative and can be misleading (*32*). A total of 662,432,037 bp (65.98% of the yellow-tufted honeyeater autosomal genome) were assigned valid GERP scores.

Proxies of individual genetic load

Because mutations occurring in evolutionarily conserved sites (as indicated by GERP scores) are assumed to be harmful, proxies of individual genetic load were obtained from the number of derived alleles present in each individual (cf. ancestral alleles).

*Ancestral allele inference.* The ancestral allele for each site was defined using the genomes of *Lichenostomus melanops cassidix* and two sister species: blue-faced honeyeater (*Entomyzon cyanotis*), and white-plumed honeyeater (*Ptilotula penicillata*; *61*). At each genomic position, the major allele among the three honeyeaters was designated as the ancestral allele of the yellow-tufted honeyeater. For sites in which not all three species had an aligned nucleotide, the ancestral allele was deemed unknown. We were able to infer the ancestral state of 767,602,866 bp (76.46% of the yellow-tufted honeyeater autosomal genome). The assignment of an ancestral allele and a valid GERP score was possible for 58.36% of the autosomal genome (585,964,369 bp).

*Proxies for individual genetic load.* Of the 26,527,283 autosomal SNPs in Dataset 2, 11,990,627 positions had a valid GERP score and an inferred ancestral allele, and could therefore be used to assess individual genetic load. With an autosomal genome size of ~1 Gb, this corresponds to an average of one SNP every ~84 bp (higher than other studies; e.g., *20* yielded one SNP per 300 bp). This 11,990,627 SNPs dataset was used to calculate all genetic load proxies described below.

A total of 28 individual genetic load proxies were applied, 20 of which were calculated following eight published studies that used GERP (Table 1). Despite the varied terminology used across these studies, all proxies could be assigned to one of three categories: realised genetic load (derived alleles in homozygous state), masked genetic load (derived alleles in heterozygous state), and total genetic load (all derived alleles regardless of their state). We termed the categories "masked" and "realised" load because most of the followed studies assumed a dominance coefficient $h = 0$ for derived alleles (i.e., complete recessiveness; except *33*) or otherwise did not explicitly state it. Five additional genetic load proxies were derived from those in the published studies to allow for downstream comparison of fitness prediction (marked with asterisks in Table 1 and Figs. 2, 3, 4, S3, S4, S5 and S6). The remaining three are new proxies we derived in this study (Table 1, figs. S3, S4, S5 and S6).

Genetic load proxies could be further categorised based on (1) whether they count the *sites* on which derived alleles occurred in an individual, or the *derived alleles* themselves (i.e., a site with a derived allele in homozygous state is counted as one in the former, and as two in the latter); and (2) how they are mathematically calculated as: (i) counts, in which derived alleles/sites in an individual are counted, (ii) sums, in which the GERP score of the derived alleles/sites in an individual are summed, and (iii) averages, in which the total sum of the GERP scores of the derived alleles/sites in an individual is standardised by a number. Classification of proxies into these categories facilitate visualization and comparison (Figs. 2, 3, 4, S3, S4, S5 and S6). The genetic load proxies were calculated following the definitions in Table 1 using custom python scripts. Because *20* selected a GERP threshold of 4 based not on its absolute value but on the distribution of GERP scores and their corresponding predicted deleteriousness (as indicated by Ensembl's Variant Effect Predictor), we approximated their rationale by identifying the percentile that a GERP score of 4 represented in their data (i.e., 92.4th percentile) and used that percentile as our threshold.

Individual inbreeding

*ROH identification*. We detected Runs of Homozygosity (ROH) using the likelihood ratio method outlined by *62*. Briefly, for 100-SNPs sliding windows and 10-SNPs step size, we calculated SNPs genotype probabilities assuming identity-by-descent (IBD) and non-IBD, while tolerating 2% heterozygous genotypes within IBD segments (to account for sequencing errors, mapping errors, and mutations). A logarithm of the odds (LOD) score was calculated for the ratio of these probabilities across all loci in the window, and windows with LOD > 0 were considered putatively IBD (*36*). Overlapping IBD windows were concatenated to form ROHs.

*Inbreeding coefficients based on ROHs ($F_{ROH}$)*. Seven different $F_{ROH}$ coefficients were calculated for each individual. $F_{ROH(3Mb+)}$, was calculated as the proportion of the genome of an individual that was in ROHs equal or longer than 3 Mb long. $F_{ROH(long)}$, $F_{ROH(medium)}$, $F_{ROH(short)}$ were calculated as the proportion of the genome of an individual that was in long (≥ 1,317,530 bp), medium (1,317,529–65,735 bp) or short ROHs (< 65,735 bp), respectively. The thresholds for short, medium and long ROHs were determined by calculating the cumulative length of all ROHs across all *cassidix* individuals and dividing it into three compartments of equal total length. $F_{ROH(recent)}$ and $F_{ROH(old)}$ were estimated as the proportion of the genome of an individual that were in ROHs that arose from recent (i.e., < 61 generations) and older (i.e., ≥ 61 generations) inbreeding, respectively. We chose 61 generations to encompass inbreeding that occurred from the start of the population contraction experienced by *cassidix* around the start of the 19$^{th}$ century (Fig. 1D). The age of ROH in generations (*g*) was calculated by adjusting a LOESS curve with span 0.2 to the linkage map of *cassidix* and interpolating each ROHs genetic map length (in cM). The number of generations (*g*) was calculated by dividing 100 by twice the interpolated genetic map length (*63*). Finally, $F_{ROH}$ was calculated as the proportion of the genome of an individual that was in ROHs of any length (i.e., ≥ 100 SNPs).

*Inbreeding coefficient based on genome-wide homozygosity ($F_{HOM}$)*. $F_{HOM}$ was calculated as the proportion of the genotyped genome of an individual in homozygous state. Unbiased estimates of genome-wide heterozygosity were obtained from Dataset 1 using *piawka*. $F_{HOM}$ was calculated as 1-$F_{HET}$.

Statistical analyses

*Fitness prediction.* We fitted generalised linear models with Poisson distribution to estimate the association of each genetic load proxy or inbreeding coefficient with LRS (42 *cassidix* individuals). Poisson-distributed GLMs have the advantage of accounting for the excess of zeros in the distribution of LRS (*64*). Genetic load proxies and inbreeding coefficients were standardized (i.e., *z*-transformed) to ensure comparability (*62*). We controlled for sex in each model (*24*), and model significance was tested with a drop-in-deviance test (*65*). Compliance with model assumptions (i.e., linearity and response distribution) was evaluated by visually inspecting model residuals, and models that violated them were considered not significant. False discovery rate due to multiple testing was controlled by adjusting *p*-values with the R function *p.adjust* (method = fdr; *66*). The same process was repeated for lifespan (43 *cassidix* individuals).

We tested whether combining inbreeding and genetic load predicted LRS better than inbreeding alone. For this, we used the realised GL proxy and inbreeding coefficient that best predicted LRS: *realised Grossen 6+* and $F_{ROH(3Mb)}$. We created a GLM with Poisson distribution including sex, $F_{ROH(3Mb)}$ and *realised Grossen 6+* as predictors of LRS. We used the R function *summary* to obtain the significance of each predictor after accounting for the effect of all other terms. The statistical effect of *realised Grossen 6+* was not significant nor did it improve the prediction of LRS ($p = 0.06$). Due to the collinearity between *realised Grossen 6+* and $F_{ROH(3Mb)}$ ($r = 0.753$, $t = 7.23$, DF = 40, *p-value* < 0.001), we also explored whether combining them into orthogonal variables would improve the prediction of LRS. We performed a partial least squares regression–designed for highly correlated independent variables–with Poisson distribution using the R package *plsRglm* (*67*), and used the Akaike Information Criterion (AIC) to choose the number of principal components to keep. The PC that maximized the prediction of LRS was used as a predictor to a final GLM model that controlled for sex. We assessed whether using the PC improved LRS prediction by comparing the $R^2$ and AIC values to the model with $F_{ROH(3Mb)}$ and sex as predictors.

*Differences between subspecies.* We tested whether there were significant differences in genetic load proxies and inbreeding coefficients between genetic groups. Subspecies *cassidix* was divided into two temporal groups (*cassidix_pre2000*: $n = 26$; and *cassidix_post2005*: $n = 17$), and *melanops* and *meltoni* were combined into group *melanops+meltoni* due to low differentiation found from our analyses. Differences among *cassidix_pre2000, cassidix_post2005*, *gippslandicus* and *melanops+meltoni* were tested using linear models, with each genetic load proxy and inbreeding coefficient as a response variable, resulting in a total of 36 models. Welch's t-tests were used for *post hoc* pairwise comparisons between groups. To account for multiple testing, we applied false discovery rate correction with the Benjamini & Hochberg method to adjust *p*-values (*66*). All models remained significant after multiple-testing correction. For easier visualisation, we plotted the mean percentage difference of each group relative to *gippslandicus* (i.e., percentage increase or decrease compared to *gippslandicus;* Figs. 4 and S5), but also presented absolute values (fig. S6).

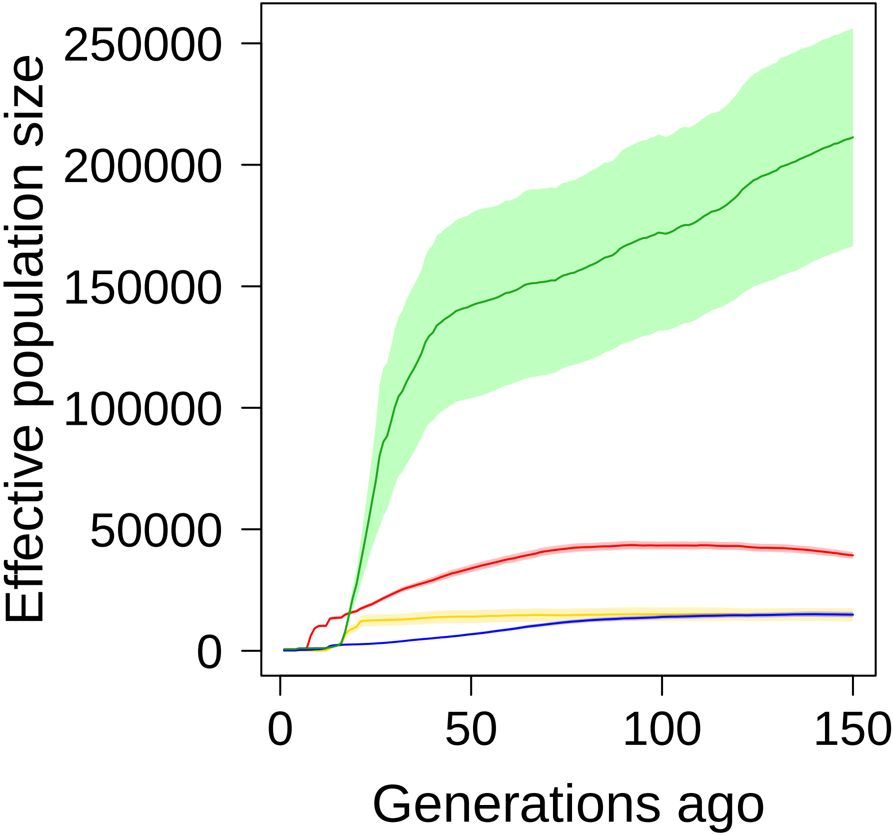

Effective population size
250000
200000
150000
100000
50000
0
0
50
100
150
Generations ago

**Fig. S1. Reconstructed demographic history of *cassidix* and *gippslandicus*.** Subspecies *cassidix* in yellow ($n$ = 13), and *gippslandicus* in red ($n$ = 17; all available samples), blue ($n$ = 13; random sample), and green ($n$ = 8; samples from a small geographical region). Shaded area represents 95% CI from 100 replicates of 700,000 SNPs each.

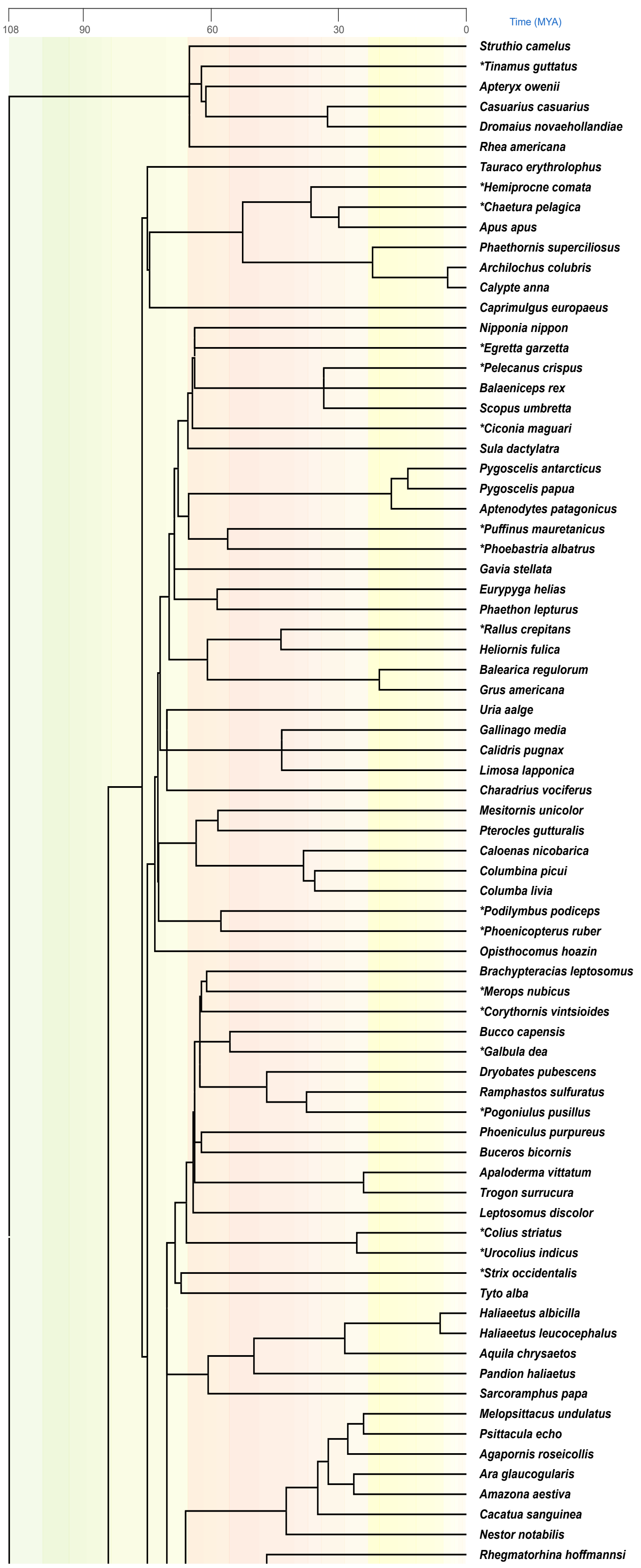

108
90
60
30
0
Time (MYA)
Struthio camelus
*Tinamus guttatus
Apteryx owenii
Casuarius casuarius
Dromaius novaehollandiae
Rhea americana
Tauraco erythrolophus
*Hemiprocne comata
*Chaetura pelagica
Apus apus
Phaethornis superciliosus
Archilochus colubris
Calypte anna
Caprimulgus europaeus
Nipponia nippon
*Egretta garzetta
*Pelecanus crispus
Balaeniceps rex
Scopus umbretta
*Ciconia maguari
Sula dactylatra
Pygoscelis antarcticus
Pygoscelis papua
Aptenodytes patagonicus
*Puffinus mauretanicus
*Phoebastria albatrus
Gavia stellata
Eurypyga helias
Phaethon lepturus
*Rallus crepitans
Heliornis fulica
Balearica regulorum
Grus americana
Uria aalge
Gallinago media
Calidris pugnax
Limosa lapponica
Charadrius vociferus
Mesitornis unicolor
Pterocles gutturalis
Caloenas nicobarica
Columbina picui
Columba livia
*Podilymbus podiceps
*Phoenicopterus ruber
Opisthocomus hoazin
Brachypteracias leptosomus
*Merops nubicus
*Corythornis vintsioides
Bucco capensis
*Galbula dea
Dryobates pubescens
Ramphastos sulfuratus
*Pogoniulus pusillus
Phoeniculus purpureus
Buceros bicornis
Apaloderma vittatum
Trogon surrucura
Leptosomus discolor
*Colius striatus
*Urocolius indicus
*Strix occidentalis
Tyto alba
Haliaeetus albicilla
Haliaeetus leucocephalus
Aquila chrysaetos
Pandion haliaetus
Sarcoramphus papa
Melopsittacus undulatus
Psittacula echo
Agapornis roseicollis
Ara glaucogularis
Amazona aestiva
Cacatua sanguinea
Nestor notabilis
Rhegmatorhina hoffmannsi

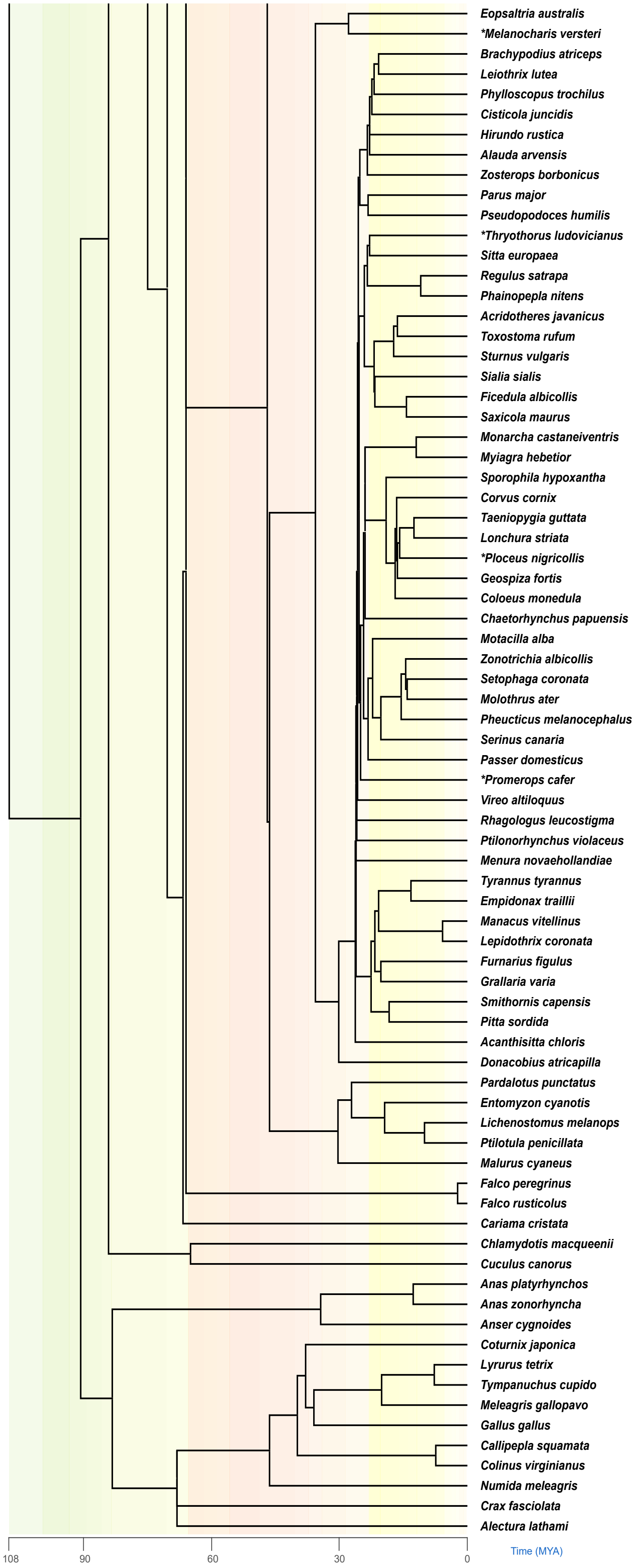
Eopsaltria australis
*Melanocharis versteri
Brachypodius atriceps
Leiothrix lutea
Phylloscopus trochilus
Cisticola juncidis
Hirundo rustica
Alauda arvensis
Zosterops borbonicus
Parus major
Pseudopodoces humilis
*Thryothorus ludovicianus
Sitta europaea
Regulus satrapa
Phainopepla nitens
Acridotheres javanicus
Toxostoma rufum
Sturnus vulgaris
Sialia sialis
Ficedula albicollis
Saxicola maurus
Monarcha castaneiventris
Myiagra hebetior
Sporophila hypoxantha
Corvus cornix
Taeniopygia guttata
Lonchura striata
*Ploceus nigricollis
Geospiza fortis
Coloeus monedula
Chaetorhynchus papuensis
Motacilla alba
Zonotrichia albicollis
Setophaga coronata
Molothrus ater
Pheucticus melanocephalus
Serinus canaria
Passer domesticus
*Promerops cafer
Vireo altiloquus
Rhagologus leucostigma
Ptilonorhynchus violaceus
Menura novaehollandiae
Tyrannus tyrannus
Empidonax traillii
Manacus vitellinus
Lepidothrix coronata
Furnarius figulus
Grallaria varia
Smithornis capensis
Pitta sordida
Acanthisitta chloris
Donacobius atricapilla
Pardalotus punctatus
Entomyzon cyanotis
Lichenostomus melanops
Ptilotula penicillata
Malurus cyaneus
Falco peregrinus
Falco rusticolus
Cariama cristata
Chlamydotis macqueenii
Cuculus canorus
Anas platyrhynchos
Anas zonorhyncha
Anser cygnoides
Coturnix japonica
Lyrurus tetrix
Tympanuchus cupido
Meleagris gallopavo
Gallus gallus
Callipepla squamata
Colinus virginianus
Numida meleagris
Crax fasciolata
Alectura lathami
108
90
60
30
0
Time (MYA)

**Fig. S2. Phylogenetic tree of 152 avian species used for GERP.**

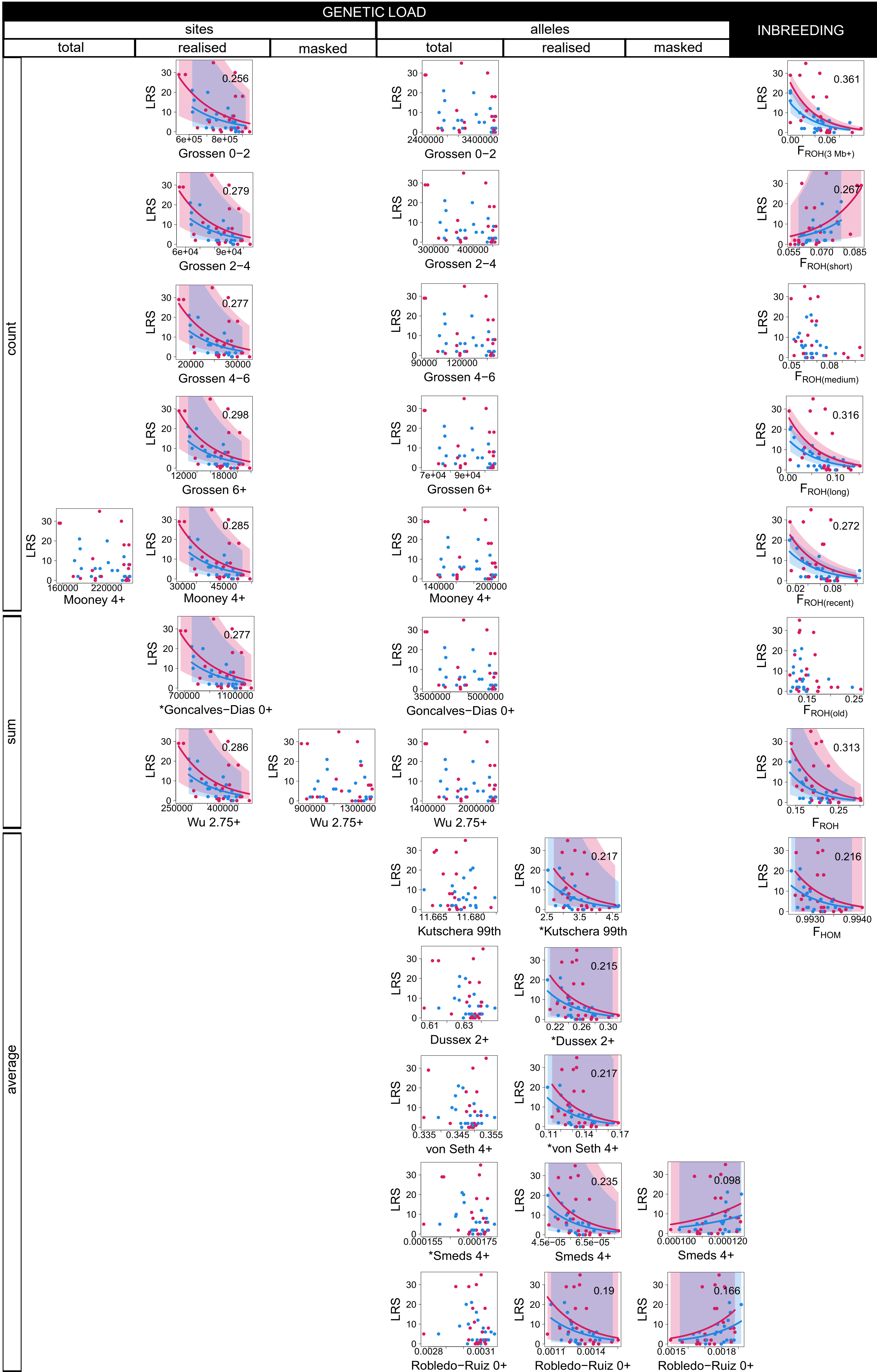
GENETIC LOAD
sites
alleles
INBREEDING
total
realised
masked
total
realised
masked
count
sum
average
LRS
0.256
Grossen 0−2
0.279
Grossen 2−4
0.277
Grossen 4−6
0.298
Grossen 6+
Mooney 4+
0.285
Mooney 4+
0.277
*Goncalves−Dias 0+
0.286
Wu 2.75+
Wu 2.75+
Grossen 0−2
Grossen 2−4
Grossen 4−6
Grossen 6+
Mooney 4+
Goncalves−Dias 0+
Wu 2.75+
Kutschera 99th
0.217
*Kutschera 99th
Dussex 2+
0.215
*Dussex 2+
von Seth 4+
0.217
*von Seth 4+
*Smeds 4+
0.235
Smeds 4+
0.098
Smeds 4+
Robledo−Ruiz 0+
0.19
Robledo−Ruiz 0+
0.166
Robledo−Ruiz 0+
0.361
F ROH(3 Mb+)
0.267
F ROH(short)
F ROH(medium)
0.316
F ROH(long)
0.272
F ROH(recent)
F ROH(old)
0.313
F ROH
0.216
F HOM

**Fig. S3. Lifetime Reproductive Success (LRS, number of fledglings produced over an individual's lifespan; y-axes) and its relationship to proxies of individual genetic load (GERP scores; x-axes) and to inbreeding coefficients (*F*; x-axes).** Genetic load proxies are grouped by columns according to whether they use counts of derived *alleles* or the *sites* in which derived alleles occurred, and whether they represent total, realised or masked load. Grouping by rows follows whether the proxies were counts of derived alleles/sites, the sum of the GERP scores for the derived alleles/sites, or average GERP scores. Names of genetic load proxies include the GERP threshold above which a mutation was considered deleterious; asterisks indicate proxies that were not calculated in the original publications but were calculated in this study for comparison purposes. Each dot represents an individual (females in red, males in blue). Trend lines represent statistically significant generalised linear models (*multiple-testing adjusted* $p < 0.001$), and shaded areas 95% CI. $R^2$ values are placed on the top right corner of statistically significant models. Only a subset of these metrics was included in Fig. 2 due to space limitations.

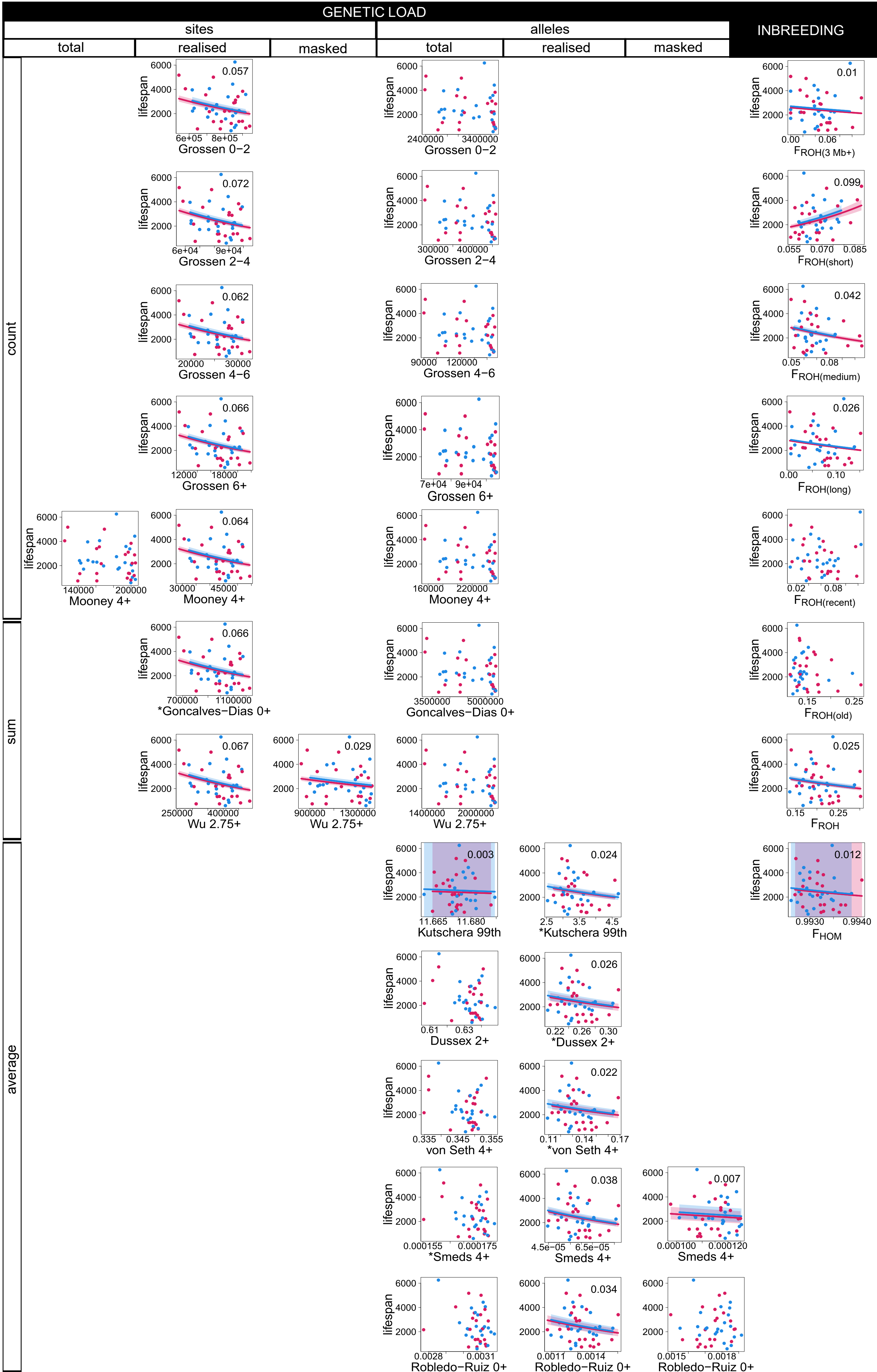
GENETIC LOAD
sites
alleles
INBREEDING
total
realised
masked
total
realised
masked
count
sum
average
lifespan
Grossen 0−2
Grossen 2−4
Grossen 4−6
Grossen 6+
Mooney 4+
*Goncalves−Dias 0+
Goncalves−Dias 0+
Wu 2.75+
Kutschera 99th
*Kutschera 99th
Dussex 2+
*Dussex 2+
von Seth 4+
*von Seth 4+
*Smeds 4+
Smeds 4+
Robledo−Ruiz 0+
FROH(3 Mb+)
FROH(short)
FROH(medium)
FROH(long)
FROH(recent)
FROH(old)
FROH
FHOM
0.057
0.072
0.062
0.066
0.064
0.066
0.067
0.029
0.003
0.024
0.026
0.022
0.038
0.007
0.034
0.01
0.099
0.042
0.026
0.025
0.012

**Fig. S4. Lifespan (number of days alive; y-axes) and its relationship with proxies of individual genetic load (GERP scores; x-axes) and to inbreeding coefficients (*F*; x-axes).** Genetic load proxies are grouped by columns according to whether they use counts of derived *alleles* or the *sites* in which derived alleles occurred, and whether they represent total, realised or masked load. Grouping by rows follows whether the proxies were counts of derived alleles/sites, the sum of the GERP scores for the derived alleles/sites, or average GERP scores. Names of genetic load proxies include the GERP threshold above which a mutation was considered deleterious; asterisks indicate proxies that were not calculated in the original publications but were calculated in this study for comparison purposes. Each dot represents an individual (females in red, males in blue). Trend lines represent statistically significant generalised linear models (*multiple-testing adjusted* $p < 0.001$), and shaded areas 95% CI. $R^2$ values are placed on the top right corner of statistically significant models. Only a subset of these metrics was included in Fig. 3 due to space limitations.

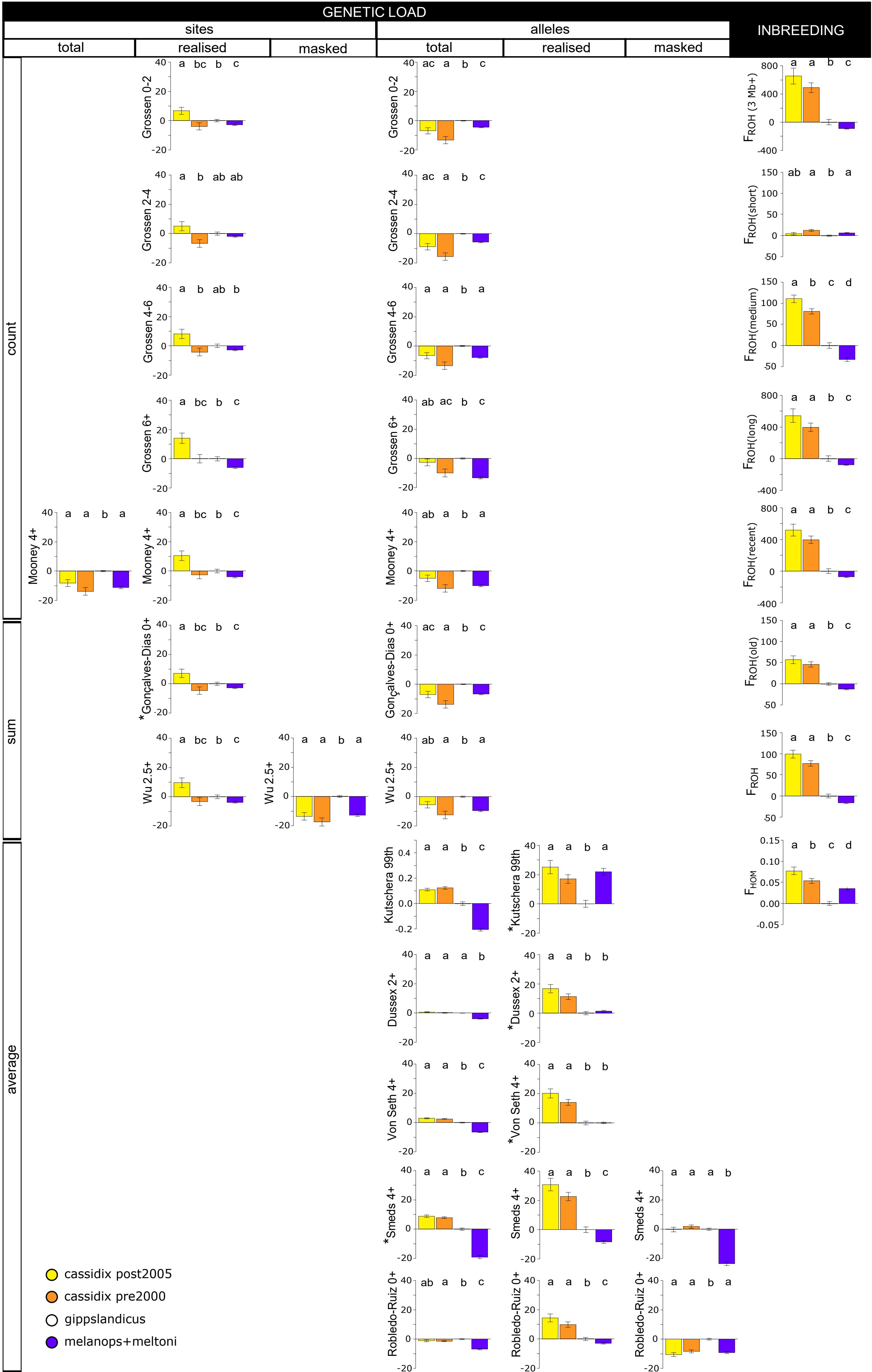
GENETIC LOAD
sites
alleles
INBREEDING
total
realised
masked
total
realised
masked
count
sum
average
Grossen 0-2
Grossen 2-4
Grossen 4-6
Grossen 6+
Mooney 4+
*Gonçalves-Dias 0+
Gonçalves-Dias 0+
Wu 2.5+
Kutschera 99th
*Kutschera 99th
Dussex 2+
*Dussex 2+
Von Seth 4+
*Von Seth 4+
*Smeds 4+
Smeds 4+
Robledo-Ruiz 0+
$F_{ROH}$ (3 Mb+)
$F_{ROH}$(short)
$F_{ROH}$(medium)
$F_{ROH}$(long)
$F_{ROH}$(recent)
$F_{ROH}$(old)
$F_{ROH}$
$F_{HOM}$
cassidix post2005
cassidix pre2000
gippslandicus
melanops+meltoni

**Fig. S5. Relative differences in proxies of individual genetic load (GERP scores) and inbreeding coefficients (*F*) between yellow-tufted honeyeater genetic groups and time periods (*cassidix post2005, cassidix pre2000*, *gippslandicus*, and *melanops+meltoni*).** Percentual differences for each group are calculated with respect to *gippslandicus*. Genetic load proxies are grouped by columns according to whether they use counts of derived *alleles* or the *sites* in which derived alleles occurred, and whether they represent total, realised or masked load. Grouping by rows follows whether the proxies were counts of derived alleles/sites, the sum of the GERP scores for the derived alleles/sites, or average GERP scores. Names of genetic load proxies include the GERP threshold above which a mutation was considered deleterious; asterisks indicate proxies that were not calculated in the original publications but were calculated in this study for comparison purposes. Bars represent mean values, and whiskers standard errors. Statically significant pairwise differences between genetic groups are represented by letters (*post hoc* Welch's t-tests, *multiple-testing adjusted* $p < 0.05$). Only a subset of these metrics was included in Fig. 4 due to space limitations.

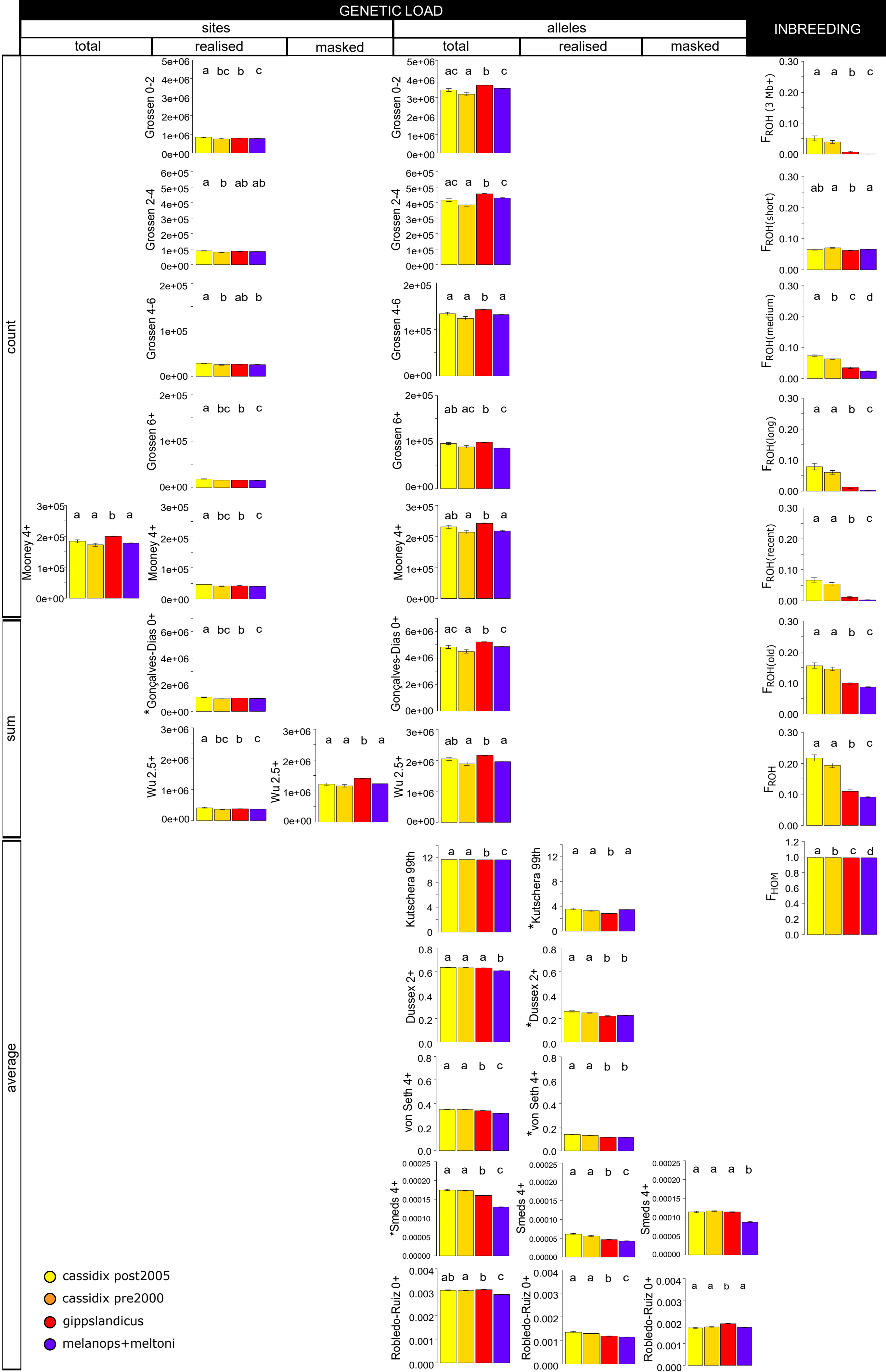

GENETIC LOAD
sites
alleles
INBREEDING
total
realised
masked
total
realised
masked
count
sum
average
Grossen 0-2
Grossen 2-4
Grossen 4-6
Grossen 6+
Mooney 4+
*Gonçalves-Dias 0+
Gonçalves-Dias 0+
Wu 2.5+
Kutschera 99th
*Kutschera 99th
Dussex 2+
*Dussex 2+
von Seth 4+
*von Seth 4+
*Smeds 4+
Smeds 4+
Robledo-Ruiz 0+
$F_{ROH}$ (3 Mb+)
$F_{ROH}$(short)
$F_{ROH}$(medium)
$F_{ROH}$(long)
$F_{ROH}$(recent)
$F_{ROH}$(old)
$F_{ROH}$
$F_{HOM}$
cassidix post2005
cassidix pre2000
gippslandicus
melanops+meltoni

**Fig. S6. Absolute values of proxies of individual genetic load (GERP scores) and inbreeding coefficients (*F*) between yellow-tufted honeyeater genetic groups and time periods (*cassidix post2005, cassidix pre2000*, *gippslandicus*, and *melanops+meltoni*).** Genetic load proxies are grouped by columns according to whether they use counts of derived *alleles* or the *sites* in which derived alleles occurred, and whether they represent total, realised or masked load. Grouping by rows follows whether the proxies were counts of derived alleles/sites, the sum of the GERP scores for the derived alleles/sites, or average GERP scores. Names of genetic load proxies include the GERP threshold above which a mutation was considered deleterious; asterisks indicate proxies that were not calculated in the original publications but were calculated in this study for comparison purposes. Bars represent mean values, and whiskers standard errors. Statically significant pairwise differences between genetic groups are represented by letters (*post hoc* Welch's t-tests, *multiple-testing adjusted* $p < 0.05$).

**Table S1. Individual data on sampling, subspecies, sex, fitness and genetic proxies of genetic load and estimates of inbreeding.**

**Table S2. List of 152 avian species used for GERP and accession numbers of their genomes.**

**Table S3. Number of species used for the phylogeny for GERP and criteria to determine ancestral alleles used by eight publications followed in this study.**
